# A novel genetic circuitry governing hypoxic metabolic flexibility, commensalism and virulence in the fungal pathogen *Candida albicans*

**DOI:** 10.1101/631960

**Authors:** Anaïs Burgain, Émilie Pic, Laura Markey, Faiza Tebbji, Carol A. Kumamoto, Adnane Sellam

**Author notes:** Corresponding authors: Prof. Adnane Sellam, Université Laval, CHU de Québec Research Center (CHUL), RC-709, 2705 Laurier Blvd, Quebec, QC, Canada G1V 4G2. and.

## Abstract

Inside the human host, the pathogenic yeast *Candida albicans* colonizes predominantly oxygen-poor niches such as the gastrointestinal and vaginal tracts, but also oxygen-rich environments such as cutaneous epithelial cells and oral mucosa. This suppleness requires an effective mechanism to reprogram reversibly the primary metabolism in response to oxygen variation. Here, we have uncovered that Snf5, a subunit of SWI/SNF chromatin remodeling complex, is a major transcriptional regulator that links oxygen status to the metabolic capacity of *C. albicans*. Snf5 and other subunits of SWI/SNF complex were required to activate genes of alternative carbon utilization and other carbohydrates related process specifically under hypoxia. *snf5* mutant exhibited an altered metabolome reflecting that SWI/SNF plays an essential role in maintaining metabolic homeostasis and carbon flux in *C. albicans* under hypoxia. Snf5 was necessary to activate the transcriptional program linked to both commensal and invasive growth. Accordingly, *snf5* was unable to maintain its growth in the stomach, the cecum and the colon of mice. *snf5* was also avirulent as it was unable to invade *Galleria* larvae or to cause damage to human enterocytes and murine macrophages. Among candidates of signaling pathways in which Snf5 might operate, phenotypic analysis revealed that mutants of Ras1-cAMP-PKA pathway, as well as mutants of Yak1 and Yck2 kinases exhibited a similar carbon flexibility phenotype as did *snf5* under hypoxia. Genetic interaction analysis indicated that the adenylate cyclase Cyr1, a key component of the Ras1-cAMP pathway, but not Ras1, interacted genetically with Snf5. Our study yielded unprecedented insight into the oxygen-sensitive regulatory circuit that control metabolic flexibility, stress, commensalism and virulence in *C. albicans*.

**Author Summary:** A critical aspect of eukaryotic cell fitness is the ability to sense and adapt to variations in oxygen concentrations in their local environment. Hypoxia leads to a substantial remodeling of cell metabolism and energy homeostasis, and thus, organisms must develop an effective regulatory mechanism to cope with oxygen depletion. *Candida albicans* is an opportunistic yeast that is the most prevalent human fungal pathogens. This yeast colonizes diverse niches inside the human host with contrasting carbon sources and oxygen concentrations. While hypoxia is the predominant conditions that *C. albicans* encounters inside most of the niches, the impact of this condition on metabolic flexibility, a major determinant of fungal virulence, was completely neglected. Here, we uncovered that the chromatin remodelling complex SWI/SNF is master regulatory circuit that links oxygen status to a broad spectrum of carbon utilization routes. Snf5 was essential for the maintenance of *C. albicans* as a commensal and also for the expression of its virulence. The oxygen-sensitive regulators identified in this work provide a framework to comprehensively understand the virulence of human fungal pathogens and represent a therapeutic value to fight fungal infections.

## Introduction

A critical aspect of eukaryotic cell fitness is the ability to sense and adapt to variations in oxygen concentrations in their local environment. Hypoxia leads to a substantial remodeling of cell metabolism and energy homeostasis, and thus, organisms must develop an effective regulatory mechanism to cope with oxygen depletion [1–3]. *Candida albicans* is an ascomycete fungus that is an important commensal and opportunistic pathogen in humans. Inside the human host, *C. albicans* colonizes predominantly oxygen-poor niches such as the gastrointestinal (GI) tract and vagina but also oxygen-rich environments such as cutaneous epithelial cells and oral mucosa [4]. This suppleness requires an effective mechanism to reprogram reversibly the metabolism to sustain the growth and to maintain the energy homeostasis. In pathogenic fungi, hypoxia impacts different virulence traits and also influences fungal fitness inside the human host [4]. For instance, isolates of the filamentous pathogenic fungus *Aspergillus fumigatus* that exhibit a higher *in vitro* fitness when oxygen is depleted were hypervirulent in an invasive model of pulmonary aspergillosis [5]. In *C. albicans*, hypoxic environment stimulates the invasive filamentous growth and the fitness inside the host [4,6,7]. Adaptation to hypoxia is also critical for the formation of the highly resistant biofilm of *C. albicans* [8,9]. Furthermore, *C. albicans* biofilms provide a hypoxic microenvironment that promotes the growth of pathogenic bacteria such as *Clostridium perfringens* [10]. Hypoxia is also an important cue that influences the host-fungal pathogens interaction. When oxygen is depleted, *C. albicans* cells mask their ß-glucans from the cell wall as a strategy to attenuate phagocytic recognition and uptake [11]. At the infection sites, *C. albicans* promotes the recruitment and the infiltration of polymorphonuclear leukocytes which consequently generate a hypoxic microenvironment that induces fungal cell masking and evasion of the immune surveillance [11].

Metabolic flexibility is essential for an opportunistic microorganism such as *C. albicans* to maintain its fitness and pathogenicity and represents thus an attractive target for antifungal therapy [12]. For instance, glyoxylate cycle (Icl1), glycolytic (Pyk1), and gluconeogenic enzymes (Pck1) are all required for the full virulence of *C. albicans* in murine model of systemic candidiasis [13]. *In vivo* studies had also demonstrated that the key glycolytic gene activator Tye7, is required for *C. albicans* to colonize the GI tract whereas Rgt1/3, a transcription factor complex that activate genes of carbon utilization, promote both GI colonization and systemic infections [14–16]. In presence of glucose, the yeast model *S. cerevisiae* induces glycolytic genes and represses genes required for the energetically demanding pathways including gluconeogenesis, glyoxylate cycles and alternative sugar utilization [13,17,18]. Unlike *S. cerevisiae*, *C. albicans* is able to activate glycolytic, gluconeogenesis and glyoxylate cycles enzymes simultaneously to utilize both glucose and alternative carbon sources [13,19]. This evolutionary advantage might help *C. albicans* to foster the efficient assimilation of complex mixtures of carbon sources to promote its fitness and virulence in the host [19]. Metabolic flexibility is also a key virulence factor in other human fungal pathogens such as *C. glabrata*, *Cryptococcus neoformans*, *A. fumigatus, Talaromyces marneffei* [20] and dermatophytes (reviewed by Ene *et al.* [13]). The therapeutic value of targeting fungal metabolism is supported by other elegant findings such as those related to manipulation of cancer metabolic flux (aerobic glycolysis) and bacterial central metabolism (folate biosynthesis) [21–23].

While hypoxia is the predominant conditions that *C. albicans* encounters inside most of the colonized human niches, the impact of this condition on metabolic flexibility has remained understudied. The transcriptional regulatory network that governs adaptation to hypoxia in *C. albicans* exhibits a high degree of complexity and interconnectedness between master transcriptional regulators such as the transcription factors Tye7, Gal4 and Ahr1 as well as the Ccr4 mRNA deacetylase [24]. Hypoxia induces a drastic remodelling of the transcriptome of *C. albicans* and other human fungal pathogens with carbohydrate transcripts including glycolysis, hexose transport, trehalose biosynthesis, fermentation and glycerol metabolism being particularly overrepresented [8,24–30]. We and other have previously shown that the transcription factor Tye7 was required for the reactivation of glycolytic genes when *C. albicans* experienced hypoxia [14,24,31]. Genetic inactivation of *TYE7* led to a significant growth defect specifically when oxygen was depleted and also to a decreased in virulence and the colonization of the GI tract [14,15]. This suggests that control of metabolism linked to the glycolytic pathway under hypoxic environments by Tye7 is a prerequisite for *C. albicans* both commensalism and infectious lifestyles. Efg1, a master transcriptional regulator of morphogenesis in *C. albicans,* was also shown to be necessary to activate glycolytic genes under hypoxia, however, this transcription factor was dispensable for the hypoxic growth [27].

So far, studies on *C. albicans* metabolism have been mainly made under normoxic conditions and undervalued the contribution of hypoxia. In this study, we performed a genetic screen to identify regulatory mechanisms that control metabolic flexibility in *C. albicans* specifically under hypoxia. This survey identified *snf5*, a mutant of a subunit of the SWI/SNF chromatin remodelling complex, with a severe growth defect specifically when utilizing alternative carbon sources under hypoxia. Snf5 was required to activate genes of alternative carbon utilization and other carbohydrate related process specifically under hypoxia. Quantitative metabolomic analysis revealed that *snf5* exhibited an altered metabolome and lipidome particularly under hypoxia demonstrating a critical role of this SWI/SNF subunit in maintaining metabolic homeostasis and carbon flux in *C. albicans*. Snf5 was also required for both commensal growth in the gut and for systemic infection suggesting that the Snf5-mediated transcriptional control of metabolic flexibility under oxygen-limiting environment is crucial for fungal fitness in the host. Our findings forge an unprecedented link between the oxygen-responsive transcriptional circuit SWI/SNF and essential functions that modulate commensalism and virulence of a human fungal pathogen.

## Results

### Survey for transcriptional regulators required for metabolic adaptation in different carbon sources under low oxygen concentration

Inside the human host, *C. albicans* colonizes predominantly oxygen-poor niches such as the GI tract but also oxygen-rich environments such as cutaneous epithelial cells and oral mucosa. This suppleness requires an effective mechanism to reprogram reversibly its metabolism in response to oxygen variation. To identify regulatory mechanisms that control carbon utilization in *C. albicans* specifically under hypoxia, mutants from the thematic transcriptional regulator libraries [32,33] were screened. A total of 836 mutant strains, representing 259 transcription regulators were screened for their ability to grow on fermentable (glucose and sucrose) and non-fermentable (glycerol) carbon sources in both hypoxia and normoxia (**Table S1**). Strains exhibiting growth defects specifically under hypoxia were considered as hit and were confirmed using the spot dilution assay. Two mutants including *tye7* and *snf5* exhibited a significant growth reduction under hypoxia (**Table S1**). Tye7p is a known transcription factor and its contribution to the growth and the re-activation of glycolytic genes under low oxygen level were already established [14]. Mutant of the SWI/SNF complex subunit, *snf5*, exhibited a severe growth defect as compared to *tye7* especially in media containing sucrose and glycerol as carbon sources. For the current investigation, we decided to focus on Snf5 as it represents a potent oxygen-dependent regulator of metabolic flexibility in *C. albicans*.

In addition to sucrose, glycerol and glucose, *snf5* growth was tested in other carbon sources including other fermentable alternative sugars (fructose, galactose, maltose), non-fermentable carbons (lactate, acetate, oleate) and polyols (sorbitol, mannitol) in both normoxia and hypoxia. No discernable growth defect was noticed for *snf5* in all tested carbon sources under normoxia (**Figure 1A**). However, under hypoxia, and in opposite to the WT and the revertant strain, *snf5* was unable to grow in media with the alternative sugar galactose, maltose, mannitol, sorbitol and the non-fermentable carbon sources lactate, acetate and oleate. As for the glucose, *snf5* growth defect was less perceptible in the presence of fructose under hypoxia. This data suggests that Snf5 is required for carbon utilization flexibility specifically in oxygen-depleted environment.

**Figure 1.**
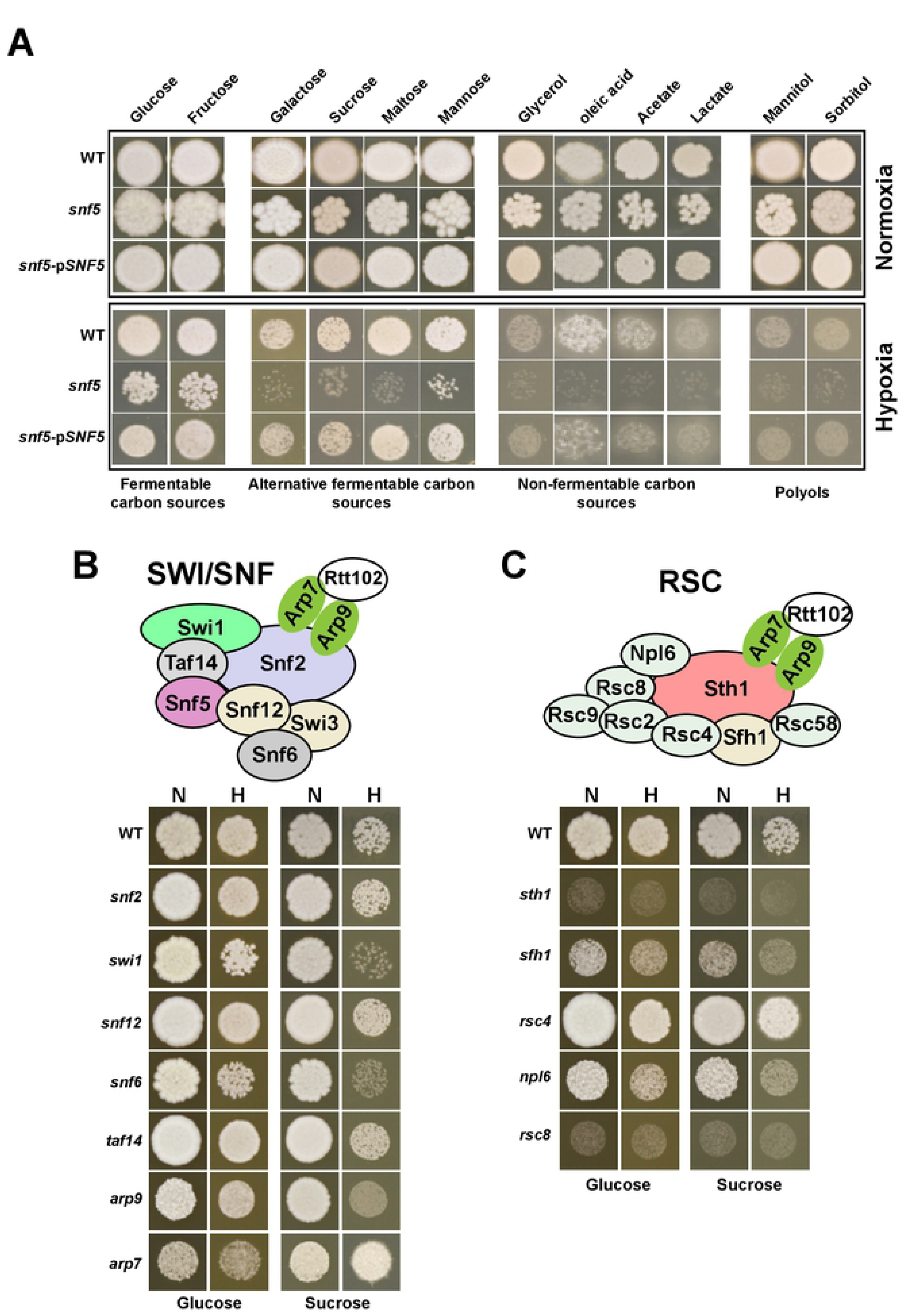
Snf5 is required for carbon metabolic flexibility specifically under hypoxia. (**A**) Overnight cultures of WT (SN250), *snf5* mutant and *snf5* strain complemented with wild type *SNF5* (*snf5*-pSNF5) cells were spotted in solid media with the indicated carbon sources under both normoxic (21% O_2_) and hypoxic (5% O_2_) conditions and incubated for 4 days at 30°C. Growth of different subunits of SWI/SNF (**B**) or RSC (**C**) chromatin remodelling complex in YP medium with either glucose or sucrose as carbon source incubated in normoxia (N) and hypoxia (H). Mutants were form the GRACE collection and were grown under repressing conditions (100 µg/ml tetracycline) for 4 days at 30°C. Schematic representation of both SWI/SNF and RSC complexes based on the architecture of their corresponding homologs in *S. cerevisiae* are shown.

### Phenotypic profiling of SWI/SNF and RSC chromatin remodeling complexes for carbon utilization under hypoxia

In addition to *snf5*, mutants of other SWI/SNF subunits including Snf12, Swi1, Arp7, Arp9, Taf14, Snf6 and the catalytic subunit Snf2 from other collections were tested for their ability to utilize glucose and sucrose under both normoxia and hypoxia. When grown on sucrose, growth defect was noticed exclusively for *swi1*, *arp9* and *snf6,* specifically under hypoxia (**Figure 1B**). No discernable growth defect was observed when those mutants were grown on glucose as a carbon source regardless the oxygen levels. When utilizing other alternative (maltose) or non-fermentable (glycerol and lactate) carbon sources, *swi1*, *snf6* and *arp9* mutants exhibited a growth defect specifically under hypoxic conditions, albeit to a lesser extent than *snf5* mutant (**Figure S1A**). These data suggest that only select subunits of the SWI/SNF complex are important for carbon metabolic flexibility under hypoxia.

We also examined the role of different SWI/SNF subunits from different fungi including the saprophytic yeast *S. cerevisiae* and the opportunist *C. glabrata* in metabolic flexibility under hypoxic environments. None of the *S. cerevisiae* mutants (*snf2*, *snf5* and *snf6*) had discernable growth defects for the tested carbon sources under either hypoxia or normoxia (**Figure S1B**). For *C. glabrata*, mutant of the catalytic subunit *snf2* exhibited growth defect in all tested sugars regardless the oxygen status while *snf6* exhibited a similar phenotype as its mutant counterpart in *C. albicans* (**Figure S1C**). On the light of these data, it is tempting to speculate that the role of SWI/SNF in metabolic flexibility is specific to fungal pathogens.

We also included to our phenotypic analysis mutant of subunits of the SWI/SNF orthologous complex, RSC, including Sth1, Sfh1, Rsc4, Rsc8 and Npl6. None of those mutants exhibited a noticeable growth defect that is oxygen-dependent (**Figure 1C**).

### Snf5 is required for the activation of genes related to carbohydrate utilization and host-pathogen interaction under hypoxia

To gain insights into cellular processes affected by the deletion of *SNF5*, we performed transcriptional profiling of both WT and *snf5* cells growing under hypoxic conditions using microarrays. YPS medium was chosen since *snf5* cells exhibited a major growth defect when using sucrose as sole carbon source under hypoxia (**Figure 1A**). We compared the transcriptional response of the WT to that of *snf5* cells and found that Snf5 was required to activate and repress 90 and 100 transcripts, respectively (**Table S2**). Among genes that Snf5 fails to activate were genes involved in sucrose metabolism including the maltase Mal32 and the maltose transporter Mal31 (**Table 1**). In *C. albicans*, sucrose utilization relies on maltase enzymes [35,36] and the lack of their inducibility might explain the growth defect of *snf5* mutant in YPS medium. Among other carbohydrate genes downregulated in *snf5*, we found the glycolytic genes *GLK1* and *GLK4* encoding both glucokinases that catalyze the phosphorylation of glucose or fructose, the first irreversible step in the intracellular metabolism of hexoses. The transcript level of genes related to carbohydrate utilization including the glucose transporter Hgt6 and Tpk2, the catalytic subunit of the cAMP-dependent protein kinase, were also downregulated in *snf5*. Repressed transcripts in *snf5* included also genes related to protein folding (Mge1, Hsp12a, Hsp12b, Asr1, Yme1), fatty acid beta-oxidation (Pex5, Pxa1, Faa2-1, Sps19), mitochondrial biogenesis (Cox16, Atp11, Coa3) and resistance to oxidative stress (Cat1, Tpk2, Yfh1) (**Table 1**). Upregulated genes were enriched in function related to ribosomal biogenesis and rRNA processing as well as amino acid and lipid biosynthetic processes and cell wall biogenesis (**Table 1**). qPCR confirmed gene expression alteration as shown by microarrays for *MAL32* and *GLK1* transcripts and also for glycolytic/neoglycogenic (*PFK1*, *FBP1*), glyoxalate (*ICL1*, *MLS1*), TCA cycle (*MDH1*) and acetate metabolism genes (*ALD6*) that were differentially regulated but did not meet the statistical filter criteria (**Figure 2B and S2, Table S3**).

**Table 1.**
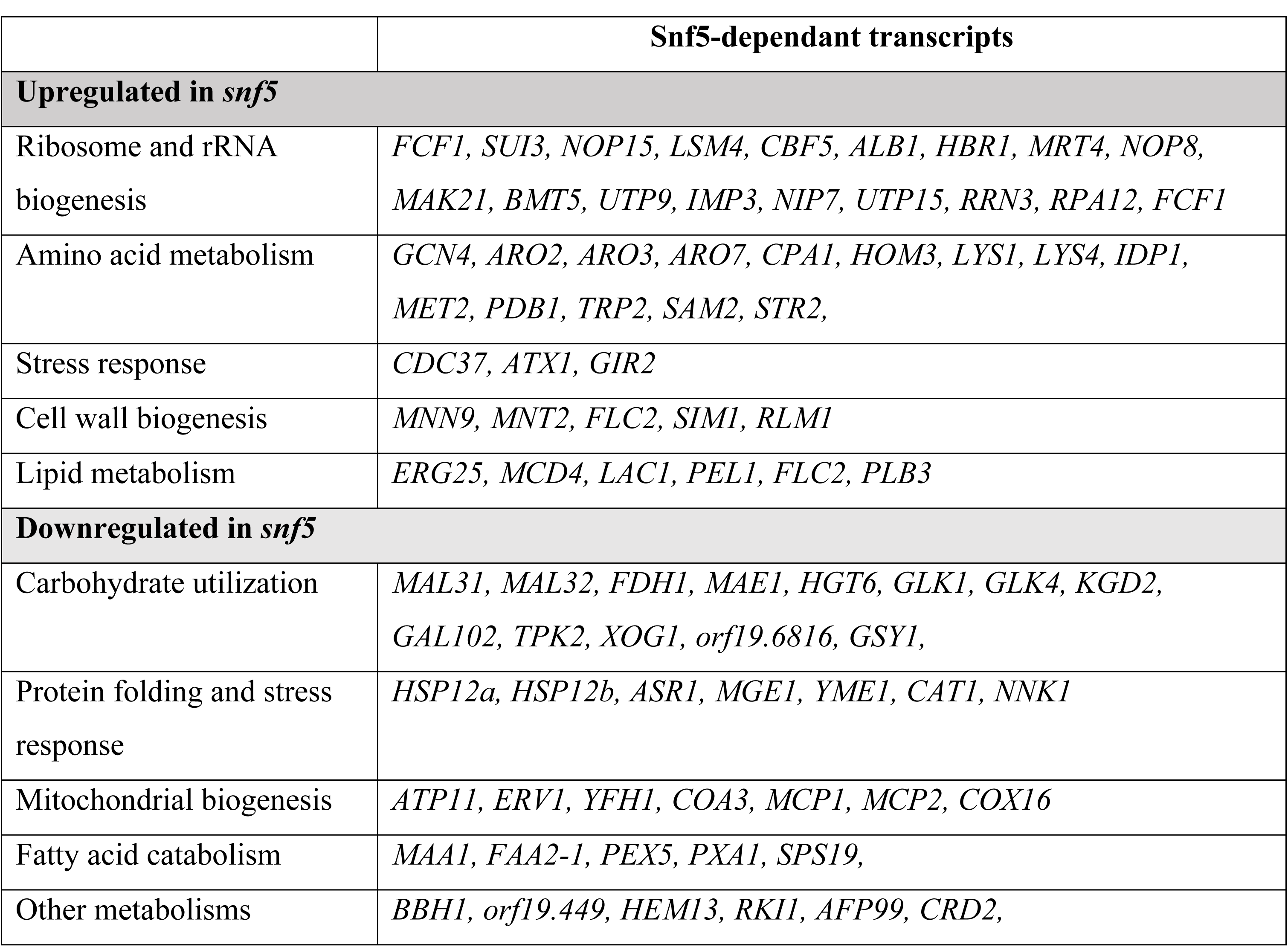
Gene function and biological process enriched in the transcriptional profile under hypoxia.

**Figure 2.**
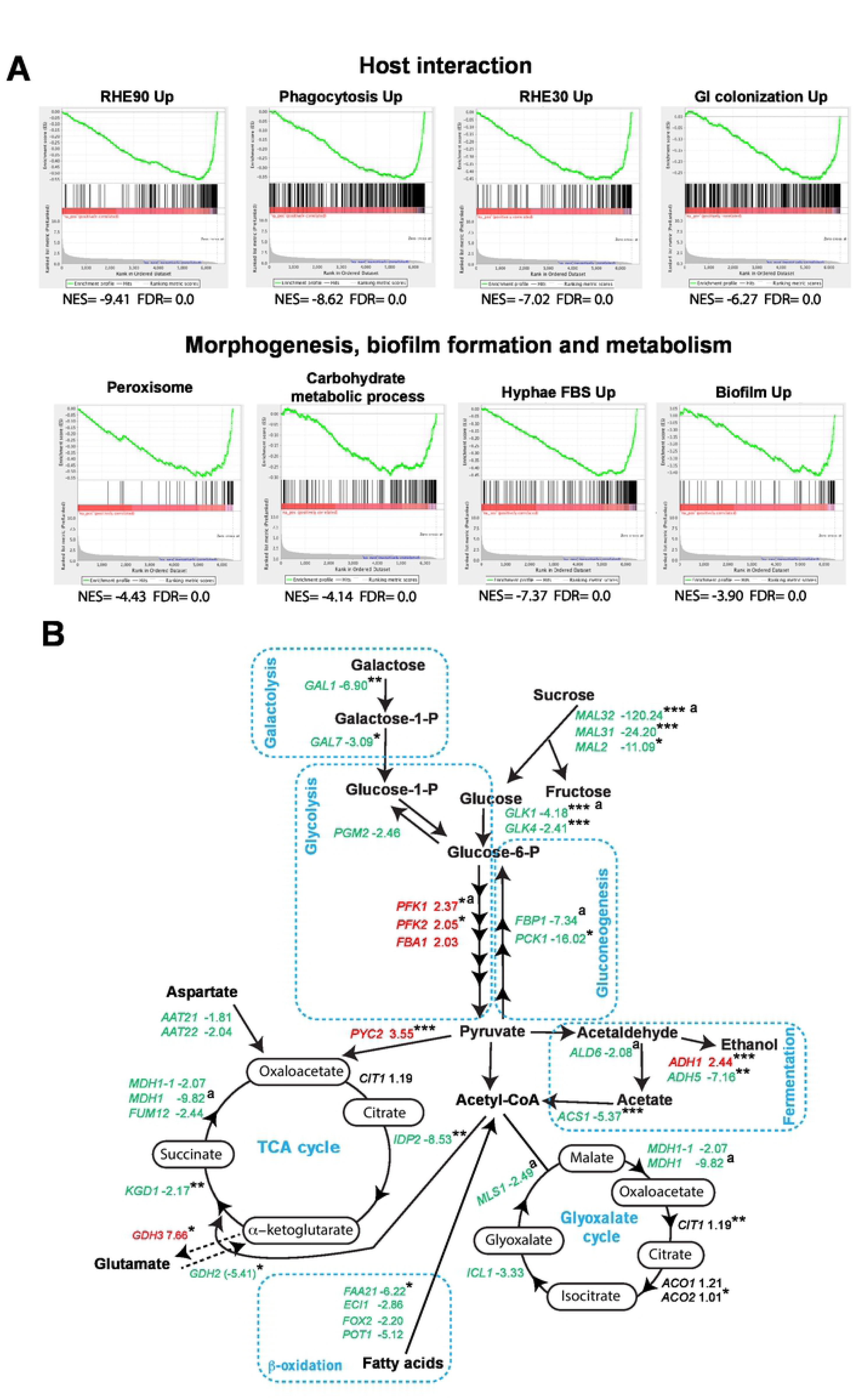
Snf5 is required for the activation of genes related to carbohydrate utilization and host-pathogen interaction under hypoxia. (**A**) Gene set enrichment analysis (GSEA) analysis of the transcriptional profile of *snf5* mutant grown in YP-sucrose under hypoxia. GSEA graphs are shown for gene sets activated during host interaction with the human oral epithelial cells infect at 30 and 90 minutes (RHE30-Up and RHE90-Up), bone marrow-derived mouse macrophages (Phagocytosis-Up) and colonization of the mammalian gut (GI colonization), or linked to virulence attributes (morphogenesis and biofilm formation) and metabolic processes (carbohydrate metabolism). NES, normalized enrichment score; FDR, false discovery rate. The complete GSEA correlations are presented in **Table S4**. (**B**) Effect of *SNF5* inactivation on central carbon metabolism of *C. albicans*. Transcript levels of genes linked to sucrose and galactose utilization, glycolysis/gluconeogenesis, fermentation, TCA and glyoxylate cycles, and beta-oxidation are shown. Upregulated and downregulated transcripts are shown in red and green, respectively. Gene with unchanged transcript level are indicated in black. P-value significance are indicated as follow: ***: p<0.05; **: p<0.08; *: p<0.1. ^a^: represents transcripts validated by qPCR in **Figure S2**.

Gene Set Enrichment Analysis (GSEA) was used to further mine the *snf5* transcriptional profile under hypoxia and determine resemblance with *C. albicans* genome annotations and other experimental large-scale omics data [24,37]. GSEA analysis confirmed the inability of *snf5* mutant to activate carbohydrate metabolic genes and fatty acid beta-oxidation (Peroxisome GO category) (**Figure 2A** and **Table S4**). Downregulated genes in *snf5* were similar to the *C. albicans* transcriptional programs expressed during the colonization of the mammalian gut [38] and the interaction with host cells including the human oral epithelial cells [39] and the bone marrow-derived mouse macrophages [40]. Furthermore, transcripts activated during the yeast-to-hyphae switch and biofilm formation were repressed in *snf5*. Taken together, these data suggest that Snf5 is a critical regulator of processes related fungal virulence traits and interaction with the host.

We have previously mapped the genomic occupancy of Snf6 using ChIP coupled to high resolution tiling arrays and showed that this SWI/SNF subunit binds directly to the promoters of genes related to carbohydrate metabolism and modulates their expression [41]. Genes associated with carbohydrate metabolism such as sucrose utilization (*MAL31*, *MAL32*), glucose and galactose metabolisms (*HGT6*, *GLK1*, *GAL7*) and TCA cycle (*KGD2*, *IDP2*) were among the direct targets of Snf6 that Snf5 fails to activate (**Figure S3**). This suggests that Snf5 is a direct regulator of aforementioned carbohydrate genes.

### Snf5 is required for transcriptional control of genes linked to the utilization of a multitude of carbon sources

To assess whether growth defect of *snf5* mutant noticed with different carbon sources, other than sucrose, correlates with gene expression misregulation, transcript levels of different genes involved in galactose (*GAL1*, *GAL7, GAL10, GAL102*), lactate (*JEN1, DLD1*) and oleic acid (*FOX2*, *POT1*, *PXA1*, *FAA21*) utilization were assessed in media with the matching carbon source under both normoxic and hypoxic conditions. For the WT strain, the transcript levels of all selected genes were significantly induced when cells experienced hypoxia (**Figure 3**). The activation of all tested genes was lost in *snf5* mutant suggesting that Snf5 is a master regulator that links oxygen status to a broad spectrum of carbon utilization routes.

**Figure 3.**
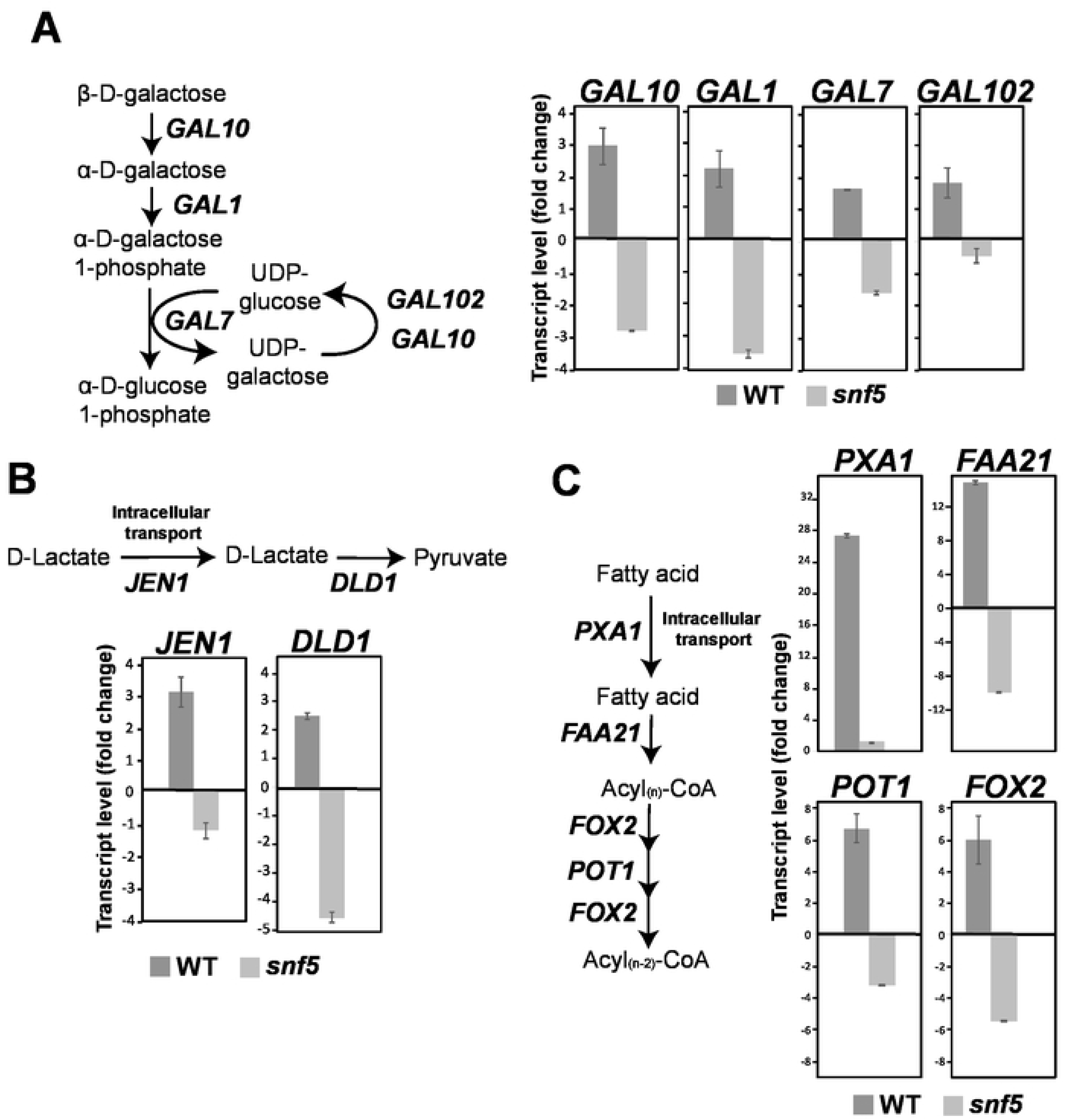
Snf5 is required for transcriptional control of genes linked to the utilization of a multitude of carbon sources. Transcript level of galactose (**A**), lactate (**B**) and oleic acid (**C**) utilization genes were measured by qPCR in the WT and *snf5* mutant strains under hypoxia relative to normoxia. Exponentially grown cells in YPD medium were washed and inoculated to fresh YP-Galactose, YP-lactate and YP-oleic acid media. Cell culture were then incubated under both normoxia and hypoxia for 1 hour at 30°C. Results are the mean of two biological replicates.

### Quantitative analysis of *snf5* metabolome uncovers a major defect of TCA cycle, beta-oxidation and Coenzyme A biosynthesis

To delineate the functional contribution of Snf5 to metabolic reprogramming of *C. albicans*, we compared the metabolome of *snf5* to that of the WT cells growing under similar conditions as used for microarray experiment with cells being exposed to hypoxia for 10 and 60 min. Consistent with the impact of *snf5* mutation in *C. albicans* metabolism under hypoxia, the PCA analysis separated the WT metabolome from that of *snf5* (**Figure 4A**). However, this analysis showed also that for either WT or *snf5* cells, the metabolome of the 10 and 60 min hypoxia was almost similar. Thus, the two hypoxic time-points were confounded for the subsequent analysis. Our metabolomic profiling uncovered that inactivation of *SNF5* affects the abundance of 389 metabolites under both normoxic and hypoxic conditions (**Figure 4B and Table S5**). Under normoxic conditions, abundance of a total of 122 metabolites was altered in *snf5* as compared to the wt. This suggests that in addition to metabolic flexibility under hypoxia, Snf5 might be crucial for *C. albicans* metabolism under normoxic conditions.

**Figure 4.**
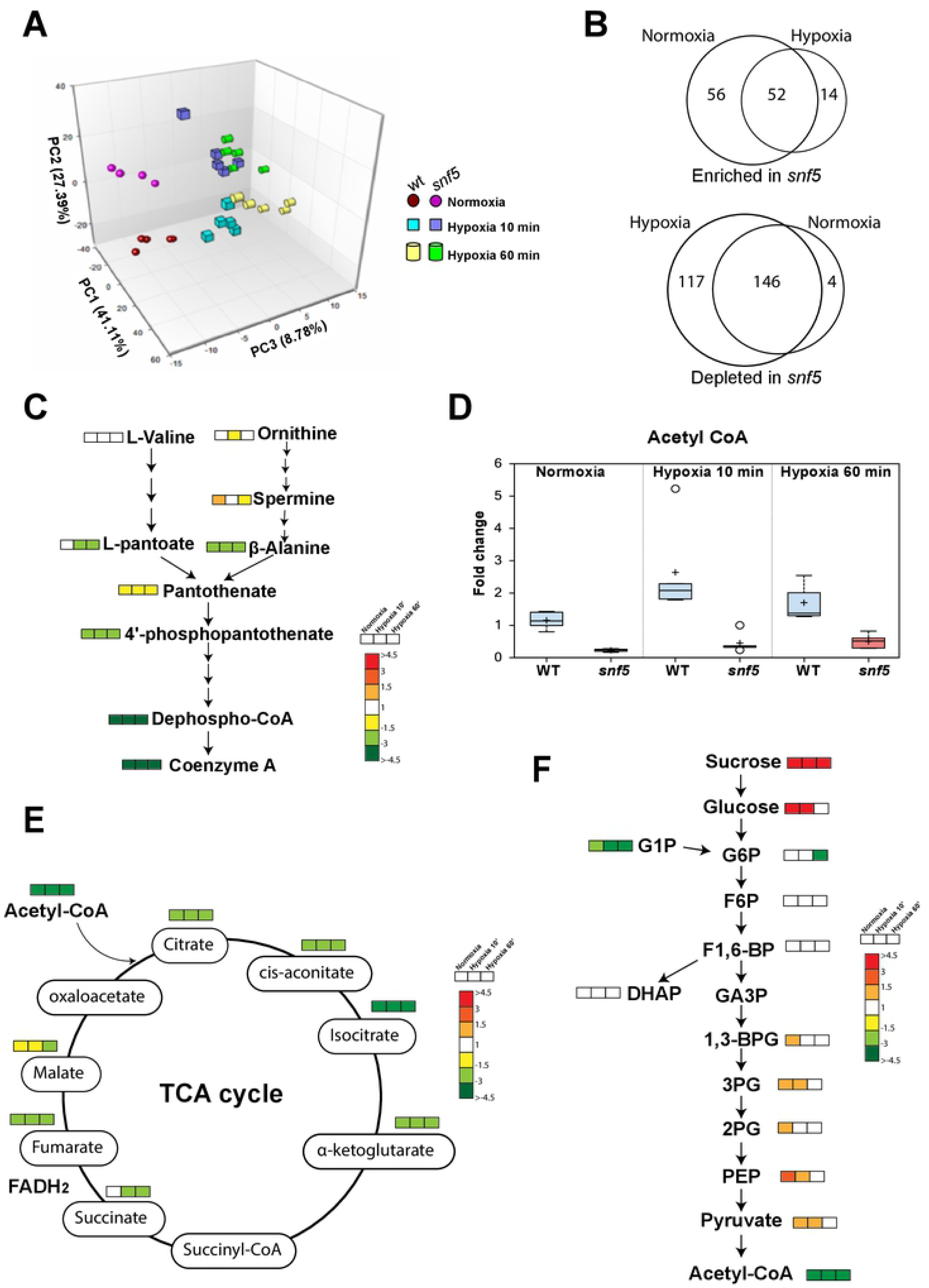
Quantitative analysis of *snf5* metabolome. (**A**) Principal component (PC) analysis of the WT (SN250) and *snf5* metabolomes exposed to hypoxia. Strains were first grown under normoxic conditions and exposed to hypoxia. Circles, squares and cylinders represent samples at 0 (normoxia), 10 and 60 min after exposure to hypoxia. PC1, PC2 and PC3 explain 41%, 27% and 8% of variance, respectively, distinguishing WT and *snf5* metabolomes. (**B**) Venn diagrams summarizing statistically significant (p<0.05) altered metabolites in *snf5* under both normoxia and hypoxia as compared to the WT strain from the total of 522, 398 and 391 detected biochemicals in samples exposed to hypoxia at 0 (normoxia), 10 and 60 min, respectively. Given that the 10 and 60 min hypoxia samples of both WT and *snf5* samples were similar, the two hypoxic time-points were confounded. (**C-F**) Summary of metabolic changes showing a major defect of Coenzyme A and acetyl CoA biosynthesis (**C-D**), tricarboxylic acid cycle (TCA) (**E**) and glycolysis (**F**).

Under both normoxic and hypoxic conditions, *snf5* mutant exhibited an increase of specific lipid classes including lysophospholipids, sphingolipids and long chain free fatty acids, while diacylglycerol lipids (DAGs), phosphatidylcholine (PCs) and phosphatidylethanolamine (PEs) were depleted (**Table S5 and S6**). A large proportion of different intermediates of purine, pyrimidine, glutathione and amino acid biosynthetic metabolism, particularly leucine and tryptophan but also phenylaniline, were elevated in *snf5* as compared to the WT (**Table S5**). *snf5* displayed also a reduced amount of pantothenate, coenzyme A and TCA cycle intermediates in addition to acetyl-CoA as compared to the WT strain regardless of the oxygen concentration (**Figure 4C-E and Table S5**). The elevated free fatty acids and the large decrease in coenzyme A biosynthesis and acetyl CoA suggest that *snf5* mutant has downregulated the oxidation of long chain FFA which is in accordance with the reduction of transcripts associated with beta-oxidation (**Figure 2B**). Similarly, the reduced level of all TCA cycle intermediates in *snf5* corroborates with the fact that many TCA cycle associated genes were down regulated in *snf5* as compared to the WT.

Under normoxic conditions*, snf5* cells had elevated baseline levels of many glycolytic intermediates, particularly glucose, PEP, and pyruvate (**Figure 4F**). This trend was also noticed when *snf5* grew under hypoxia for 10 min and disappeared after 60 min. This suggests that *snf5* cells rely more on glycolysis than WT cells under normoxic and at early exposure to hypoxia and might reflect a compensatory mechanism to circumvent the defect of sucrose utilization. Accordingly, our transcriptional profiling and qPCR experiments showed up regulation of many glycolytic genes such as *PFK1*, *PFK2* and *FBA1* together with their transcriptional regulator Tye7 (**Figure S2 and Table S3**). The depletion of acetyl CoA in *snf5* cells is most likely related to the decrease amount and biosynthetic rate of CoA. This could be explained also by the reduced transcript level of the aldehyde dehydrogenase Ald6 and the Acetyl-CoA synthetase Acs1 acting in the so-called the pyruvate dehydrogenase (PDH) bypass that convert pyruvate to CoA (**Figure 2B**). Sucrose was also elevated in *snf5* cells under both normoxia and hypoxia reflecting their inability to metabolize this disaccharide (**Figure 4F**).

Most of the depleted metabolites in *snf5* under normoxia were also reduced under hypoxia. However, under hypoxia, a total of 117 metabolites were specifically decreased (**Figure 4B and Table S3**). This set of metabolites contains monohydroxy and long chain saturated fatty acids, and intermediates of amino acid biosynthesis including glutamine, tyrosine and leucine. Other *snf5*-depleted metabolites under hypoxia were additional metabolic intermediates that belong to the same metabolic classes that were commonly depleted in normoxia including DAGs, PCs, and different intermediates of purine and pyrimidine metabolisms (**Table S3**). In accordance with the reduced transcript level of the glucokinases Glk1 and Glk2 in *snf5*, glucose-6-phosphate amount was significantly lower specifically under hypoxia. Taken together, the alteration of *snf5* metabolome suggests that Snf5 is required for metabolic homeostasis to accommodate *C. albicans* to the metabolic demand accompanying oxygen depletion.

### Snf5 is required for adherence to enterocyte and intestinal colonization

The ability of *C. albicans* to colonize intestinal tract is fundamentally associated to its metabolic flexibility to utilize a wide range of nutrients brought directly by the daily diet or from diet by-products after being processed by the intestinal microbiota. Given the fact that hypoxia is the predominant condition in the gut, the role of Snf5 in metabolic flexibility might have a pivotal impact on GI colonization by *C. albicans*. This hypothesis is supported by the fact that *snf5* mutant was unable to activate the transcriptional program associated with the colonization of the mammalian gut (**Figure 2A**). To assess whether the colonization of GI tract by *C. albicans* requires Snf5, we used murine model where antibiotic-treated mice were orally inoculated by WT, *snf5* and the revertant strains. For each strain, intestine colonization was followed after 1, 3- and 9-days post-inoculation by determining the CFU from fresh fecal pellets. Both WT and complemented strains exhibited a similar and a sustained colonization profile across the time course while *snf5* mutant exhibited a significantly reduced degree of intestinal occupancy (**Figure 5A**). At day 10, the mice were sacrificed, and the degree of colonization was measured in the stomach, the cecum and the colon. The obtained data showed that *snf5* exhibited a reduced level of colonization in all tested organs as compared to the WT and revertant strains (**Figure 5B-D**). Since *snf5* levels were mostly below the detection threshold (1 CFU), we used qPCR to reassess its depletion from the cecum. The qPCR data recapitulated the previous finding and confirmed the colonization defect of *snf5* mutant in the cecum (**Figure 5E**).

**Figure 5.**
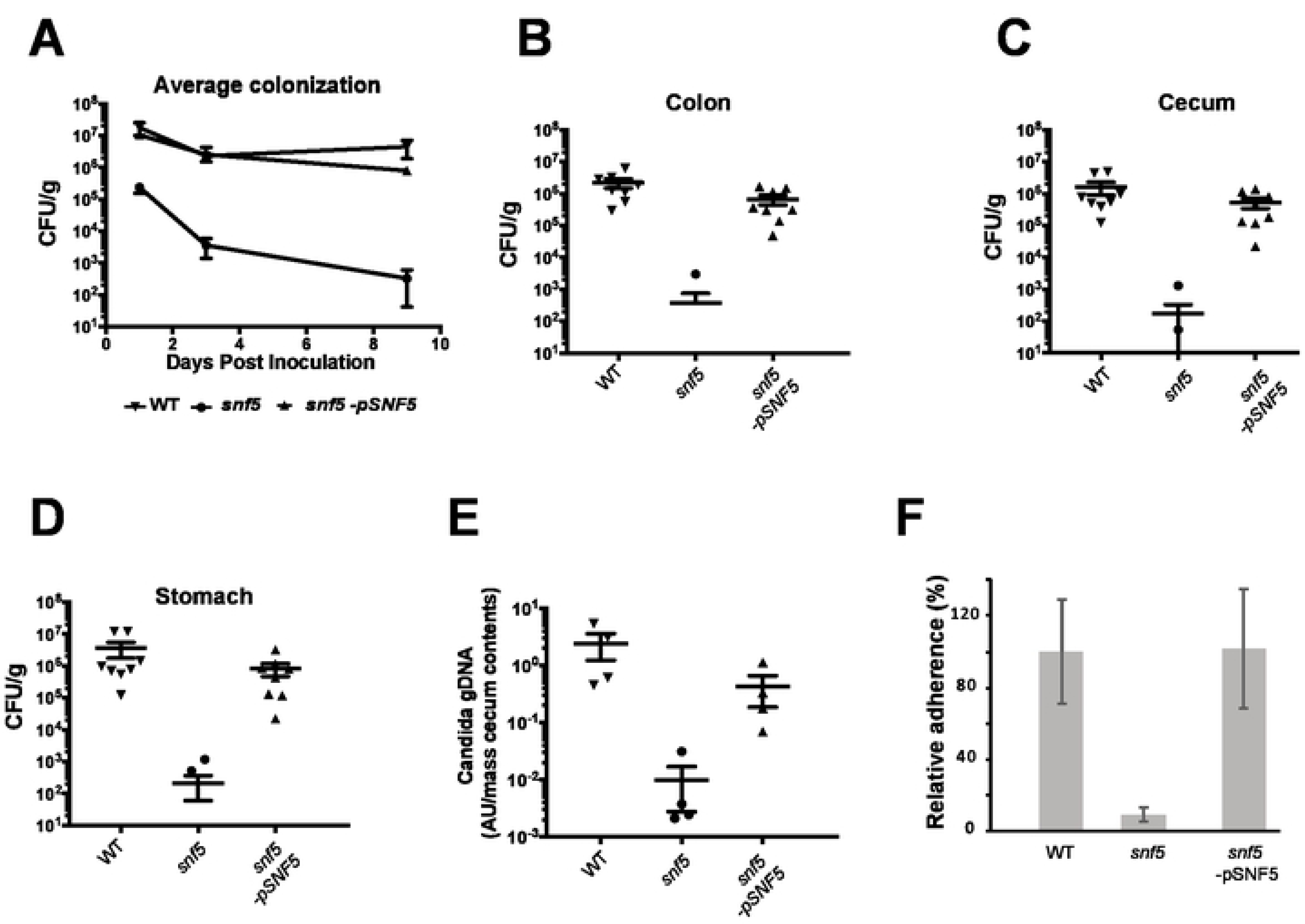
Snf5 is required for adherence to human enterocytes and intestinal colonization. Antibiotic-treated mice were orally inoculated by WT, *snf5* and the complemented strain. (**A**) Intestine colonization was followed after 1, 3- and 9-days post-inoculation by determining the CFU from fresh fecal pellets. At day 10, the mice were sacrificed, and the degree of colonization was measured in the stomach (**B**), the cecum (**C**) and the colon (**D**). Multiple means were compared using Dunn’s multiple comparison test and *snf5* colonization defect is statistically significant as compared to the WT or the complemented strain. (**E**) qPCR validation of the colonization defect of *snf5* in the cecum. (**F**) Adherence of the WT, *snf5* and the revertant strains to the HT-29 enterocytes. The adhesion was calculated as % of cells that remain attached to HT-29 cells after PBS rinsing relative to the *C. albicans* WT strain. The presented results are from at least three independent experiments performed in triplicate.

Previous work by Mitchell and collaborators had shown that Snf5 is a key regulator of adherence of *C. albicans* to biotic substrate [42]. This led us to investigate whether the failure of *snf5* to colonize GI tract might be related to defect in adhesion to intestinal host cells. Adherence of the WT, *snf5* and the revertant strains to the HT-29 enterocytes was assessed and the obtained data revealed a significant reduction of ∼ 92% in *snf5* as compared to the WT and the revertant strains (**Figure 5F**). These findings suggest that the failure of *snf5* to sustain its commensal growth in the GI tract might be a dual consequence of its inability to utilize different carbon sources and its inability to attach to the intestinal epithelial cells.

### Snf5 is required for host invasion and hyphae formation

GSEA analysis uncovered that *snf5* was unable to properly control transcripts associated with the morphogenetic switch in response to different cues including Spider and fetal bovine serum (FBS). This finding is in accordance with previous study reporting the defect of *snf5* in differentiating hyphae in liquid Spider medium at 37°C [42]. Given the oxygen-dependent role of Snf5 in carbon utilization, we tested the role of Snf5 in mediating the yeast-to-hyphae transition in response to hypoxia. While WT and the revertant strains exhibited an enhanced filamentation growth under hypoxia as compared to normoxia, *snf5* mutant grew as a smooth colony and their hyphal growth was completely compromised (**Figure 6A**). We also found that *snf5* was unable to form invasive hyphae in response to other cues including 10% FBS, N-acetylglucosamine (GlcNAc) and Spider (**Figure 6A**). Consistent with the filamentation defect, *snf5* failed to invade agar surfaces and was easily washed out from agar plate as compared to WT and the complemented strain (**Figure 6B and 6C**).

**Figure 6.**
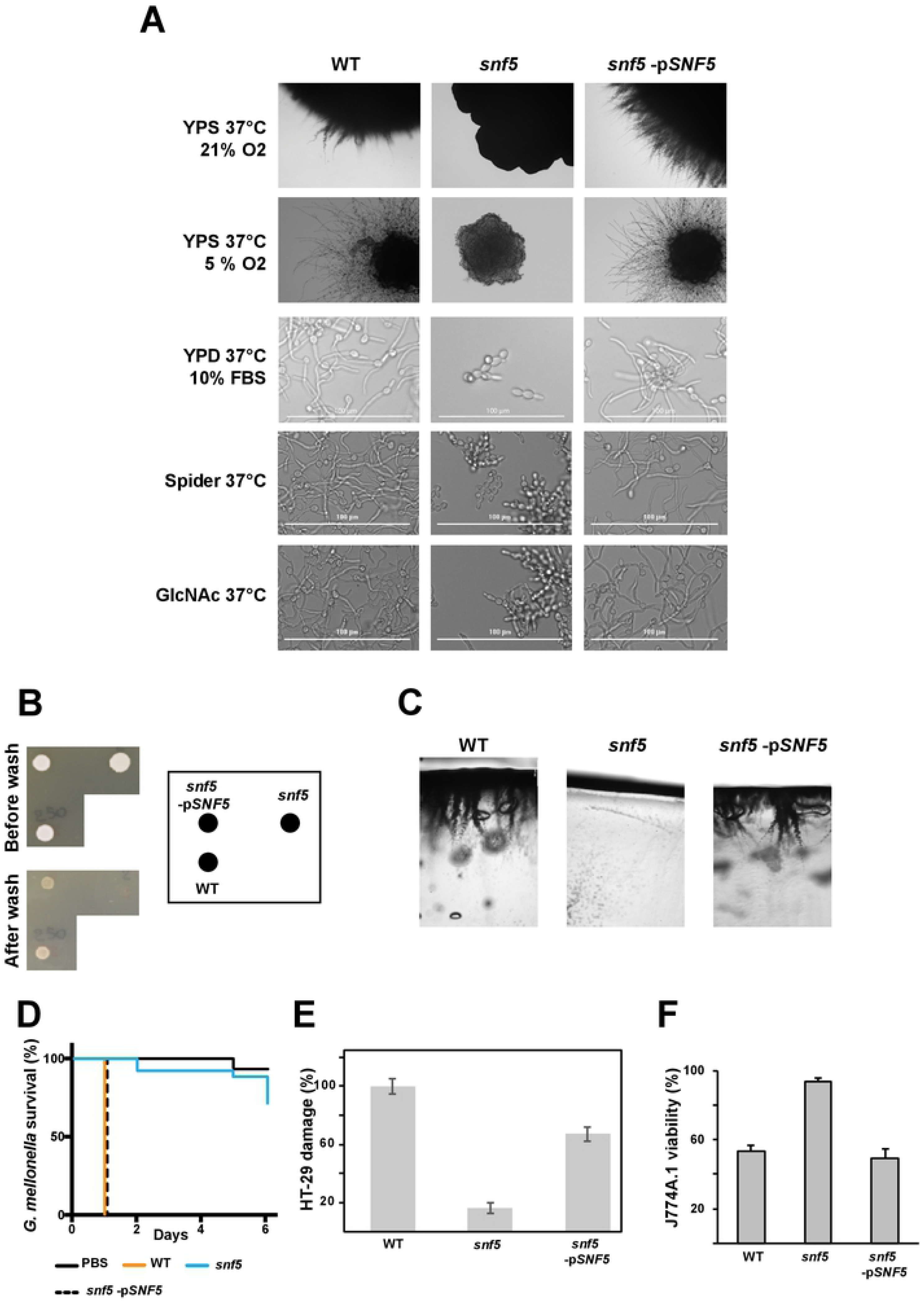
Snf5 is required for host invasion and hyphae formation in response to different cues. (**A**) Colony morphologies of the WT, *snf5* and the complemented strain after four days at 37°C on solid YPS medium under both normoxia (21% O_2_) and hypoxia (5% O_2_), or growing in in liquid Spider, and YPD media supplemented with either 10% fetal bovine serum (FBS) or 2.5 mM N-acetyl-D-glucosamine (GlcNAc) for 3 hours. (**B-C**) Agar invasion assay. Exponentially growing WT, *snf5* and revertant strains were seeded on YPD agar plate and incubated for 3 days at 30 °C. (**B**) The plates were photographed before and after being washed with PBS. (**C**) Agar sections were also performed and photographed. (**D**) *snf5* has reduced virulence in the *G. mellonella*-*C. albicans* model of systemic infection. WT, *snf5* and the revertant strains and control PBS were injected to *G. mellonella* larvae and survival was monitored daily for a period of 6 days. *SNF5* inactivation attenuates damage of the human colon epithelial HT-29 cells (**E**) and the murine macrophages J774A.1 (**F**). HT-29 cell damage was assessed using the lactate dehydrogenase (LDH) release assay and was calculated as percentage of LDH activity in cell infected by *snf5* and the revertant strains relatively to cells infected by the WT (SN250) strain. Macrophage viability was calculated as percentage of LDH activity in cell infected by the WT and the revertant strains relative to J774A.1 infected by the *snf5* strain. Results are the mean of three independent replicates.

As defect in hyphal growth is not always associated with loss of virulence [43], we used the *Galleria mellonella*-*C. albicans* model of systemic candidiasis to assess the role of Snf5 in mediating systemic infection. Injection of PBS and *snf5* mutant resulted in a death of only 5 % and 10 %, respectively, at day 5, while *G. mellonella* larvae infected by the WT or the revertant strain were completely killed at day 1 (**Figure 6D**). We also tested whether inactivation of *SNF5* led to a reduction of damage to the human colon epithelial HT-29 cells using the LDH release assay. HT-29 damage was reduced by 80% for *snf5* as compared to the WT and the revertant strain (**Figure 6E**).

*C. albicans* cells engulfed by macrophages experience nutrient deprivation and also hypoxic environment [44]. Given the role of Snf5 in mediating metabolic flexibility under hypoxia, we tested the requirement of Snf5 for the *C. albicans* survival during the interaction with the murine macrophages J774A.1. While the WT and *snf5* complemented strains were able to escape and damage 50% of J774A.1 macrophages, *snf5* mutant was inoffensive and led to 95% survival of phagocytic cells (**Figure 6F**). Taken together, in addition to the requirement of Snf5 for the intestinal commensal growth, our data revealed that this SWI/SNF subunit is also essential for *C. albicans* virulence.

### Ras1-cAMP pathway and, the Yak1 and Yck2 kinases are required for carbon metabolic flexibility under hypoxia

The GSEA analysis uncovered that the transcriptional profile of *snf5* was significantly similar to that of mutants of different signaling pathways including Ras1-cAMP-PKA (*ras1*, *cyr1*) and AMPK/Snf1 (*sak1*), as well as mutant of the serine-threonine protein kinase Yak1, the AGC family protein kinase, Sch9 and the serine-threonine phosphatase, Sit4 (**Figure 7A**). The metabolic flexibility of these mutants was tested in different carbon sources including glucose, sucrose and galactose in both normoxia and hypoxia. A *cyr1* mutant exhibited a similar growth defect to that of *snf5* when utilizing sucrose and galactose under hypoxia while *ras1*, *yak1* and *yck2* mutants showed only a moderate defect under the same conditions (**Figure 7B**). The other tested mutant including *tpk1, tpk2*, *sch9, ccr4, pop2, med31, cup2, sit4, cwt1, zap1, rgt1, rgt3,* and *sko1* had no discernable growth defect under all tested conditions. This suggests that in addition to SWI/SNF complex, the cAMP-dependent protein signalling pathway and to a lesser extent the Yck2 and Yak1 contribute to metabolic flexibility under hypoxic conditions. Intriguingly, phenotype of mutants of the two catalytic protein kinase A (PKA) subunits, Tpk1 and Tpk2 acting downstream Cyr1 [46] in media with different carbon sources under hypoxia was not distinguishable from the WT strain. This suggests that control of hypoxic metabolic flexibility in *C. albicans* by Cyr1 is PKA-independent.

**Figure 7.**
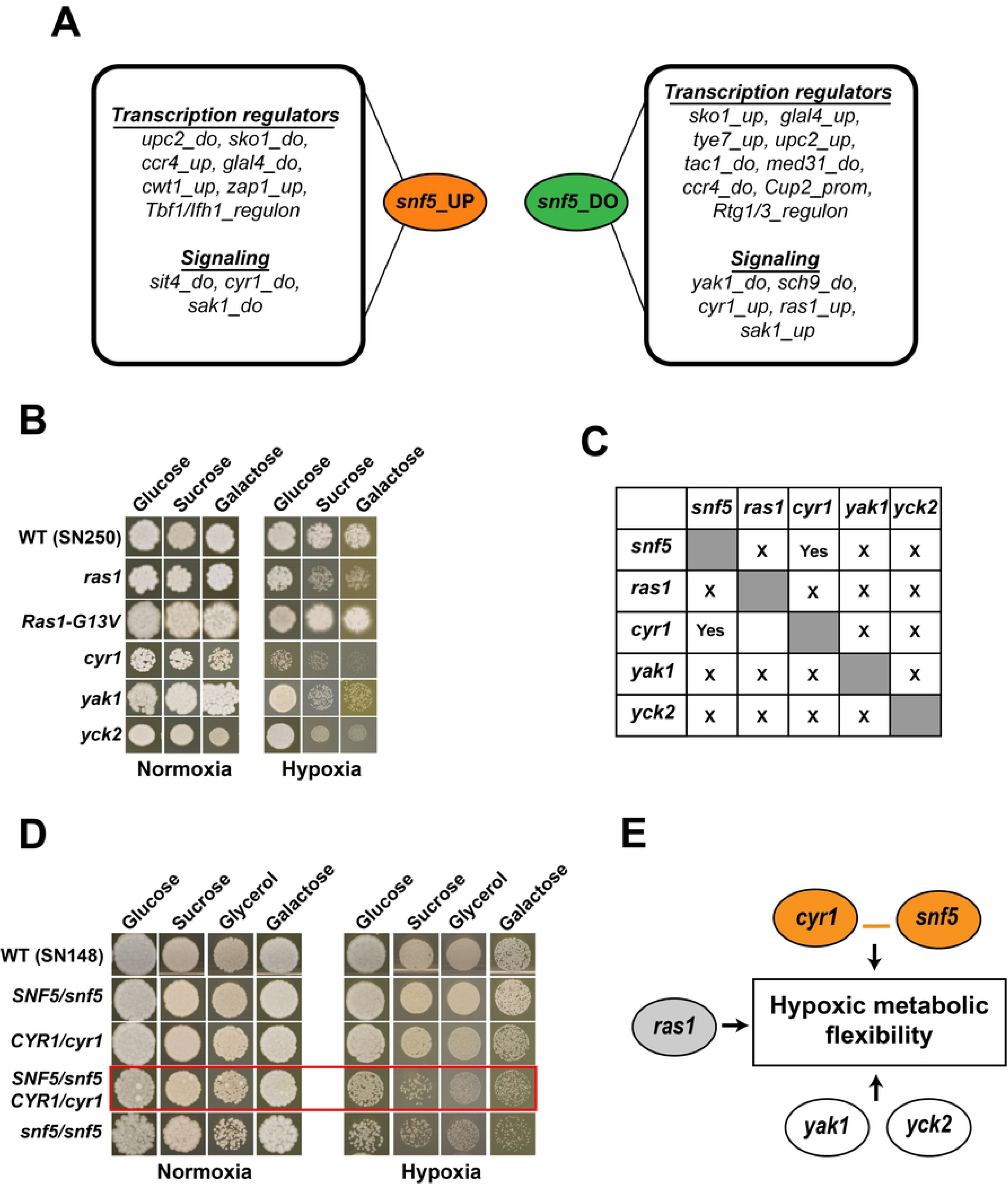
Additional signaling pathways are required for carbon metabolic flexibility under hypoxia. (**A**) Summary of GSEA analysis of up- and down-regulated transcript in *snf5* displaying correlations with *C. albicans* mutants of transcriptional regulators and signalling proteins. “Up” and “do” correspond to up- and down-regulated transcripts in a gene list, respectively. “Regulon” indicates promoter lists occupied by a transcription factor as shown by ChIP-seq or ChIP-chip. “Prom” corresponds to promoters that have the predicted cis-regulatory binding motif of a transcription factor. (**B**) Ras1-cAMP, Yck2 and Yak1 are required for carbon metabolic flexibility under hypoxia. Cultures of the WT (SN250), *yck2*, *yak1*, *cyr1*, *ras1* and the constitutively active Ras1-G13V strains were spotted in solid media with the indicated carbon source under both normoxia (21% O_2_) and hypoxia (5% O_2_) and incubated for 4 days at 30°C. (**C**) *snf5*/*SNF5* heterozygous mutant strain were tested for complex haploinsufficiency (CHI) with *yck2/YCK2*, *yak1/YAK1*, *cyr1/CYR1*, *ras1/RAS1* heterozygous mutants. Pairwise genetic interactions: “Yes”, indicates CHI interaction while “X” refers to the absence of CHI. (**D**) CHI-based genetic interaction analysis reveals a functional link between the adenylyl cyclase Cyr1 and Snf5. Normoxic and hypoxic growth of the WT (SN250), *snf5*, the *cyr1/CYR1* and *snf5*/*SNF5* heterozygous mutants, and the *cyr1/CYR1 snf5*/*SNF5* double heterozygous strain are shown in the presence of different sugars. (**E**) Summary the genetic circuit controlling the hypoxic metabolic flexibility in *C. albicans*.

### Complex haploinsufficiency-based genetic interaction analysis reveals a functional link between the adenylyl cyclase Cyr1 and Snf5

Complex haploinsufficiency (CHI) is a powerful tool to assess functional relationships between genes, and in particular, whether sets of genes function in the same or parallel pathways [47,48]. To elucidate functional connections between *SNF5* and genes of signaling pathways required for the hypoxic metabolic flexibility identified above (*RAS1*, *CYR1*, *YAK1*, *YCK2*), we used CHI concept to identify potential genetic interactions. A total of 10 double heterozygous mutants were generated as indicated in **Figure 7C** and their ability to grow in different carbon sources (glucose, sucrose and galactose) under both normoxic and hypoxic conditions was evaluated together with their parental single heterozygous strains. While both *SNF5*/*snf5* and *CYR1*/*cyr1* mutants had no perceptible fitness defect, the *SNF5*/*snf5 CYR1*/*cyr1* strain exhibited a growth defect similar to that of *snf5*/*snf5,* suggesting that Snf5 and the adenylyl cyclase Cyr1 operate on the same pathway (**Figure 7D**). Intriguingly, even if Cyr1 is a key component and an effector of Ras1 pathway [49], *SNF5* did not interact genetically with *RAS1*. Previous study had shown that *C. albicans* filamentation in response to carbon dioxide depends on Cyr1, but not on Ras1, suggesting that Cyr1 controls specific functions that are independent from the Ras pathway [50,51]. Neither *YAK1* nor *YCK2* exhibited CHI interaction with *SNF5*. Our CHI data did not support functional relatedness between *CYR1* and *YCK2*, *RAS1* or *YAK1* (**Figure 7C**). This finding suggests that Cyr1-Snf5 regulatory axis together with Ras1, Yck2 and Yak1 parallelly control metabolic flexibility in *C. albicans* under hypoxic environment.

### Snf5 is required to maintain ATP homeostasis associated with carbon metabolic flexibility under hypoxia independently from the AMP-activated protein kinase Snf1

The AMP-activated protein kinase (AMPK) Snf1 plays an important role in energy homeostasis by acting as an ATP sensor that is activated by phosphorylation following energy depletion [52]. Activated Snf1 stimulates processes leading to the production of ATP and the shut-down of ATP-consuming processes [52]. In metazoans, hypoxia reduces ATP production by lowering the activity of the electron transport chain which led to the activation of the Snf1/AMPK protein kinase [1]. Accordingly, we hypothesized that in *C. albicans*, Snf1/AMPK might be activated in response to ATP drop that accompanies oxygen depletion. This is supported by our GSEA analysis that indicated a significant similarity between the transcriptional profile of *snf5* and mutant of the kinase Sak1 that regulates the activity of Snf1 [53] (**Figure 7A**). Furthermore, in *S. cerevisiae*, Cyr1 is phosphorylated by the Snf1 to promote ATP homeostasis [54], a mechanism that might be conserved in *C. albicans*. Together, this led to hypothesize that in *C. albicans*, Snf1 might be activated under hypoxia to accommodate the metabolic demand and to promote the flexibility of carbon utilization through Cyr1-Snf5 axis.

We first measured ATP abundance in both normoxic and hypoxic conditions when *C. albicans* grew on media with glucose, sucrose and glycerol as sole source of carbon for the WT, *snf5* and the revertant strains. In the presence of glucose, both WT, *snf5* and the complemented strains had a similar amount of ATP which dropped by 12 times when shifted from normoxia to hypoxia (**Figure 8A**). Thus, as in metazoans, hypoxia led to a decrease of ATP in *C. albicans* cells. On sucrose- or glycerol-containing media, and under both normoxia and hypoxia, *snf5* exhibited a reduced ATP amount as compared to the WT (4-fold reduction under normoxia for both carbon sources and, 11- and 3-fold reduction under hypoxia for sucrose and glycerol, respectively) (**Figure 8A**). This suggests that Snf5 is required to maintain cellular ATP homeostasis regardless of oxygen levels and that the usual drop of ATP when cells experience hypoxia might exaggerate the energy deficiency phenotype of *snf5* which might led to the observed growth defect especially when utilizing sucrose and glycerol (**Figure 1A**).

**Figure 8.**
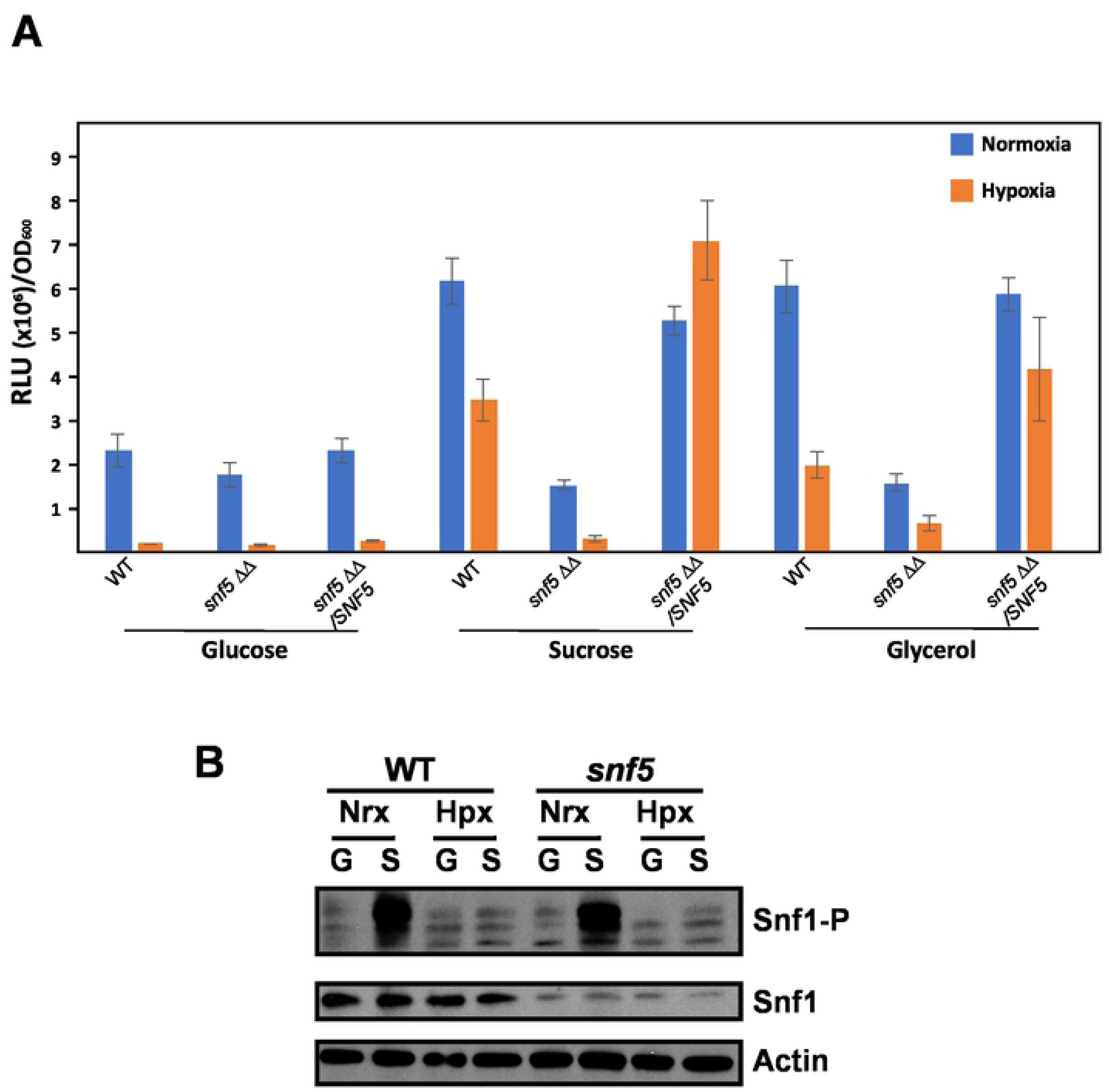
Snf5 is required to maintain ATP homeostasis associated with metabolic flexibility under hypoxia independently from the AMP-activated protein kinase Snf1. (**A**) Effect of hypoxia on cellular levels of ATP. ATP was quantified in the WT (SN250), *snf5* mutant and the complemented strains by luminescence using the BacTiter-Glo reagent in liquid medium with different carbon sources (glucose, sucrose and glycerol) and oxygen status (21 and 5 % O2). The presented results are from at least three independent experiments performed in triplicate. (**B**) Effect of hypoxia and carbon source quality on the phosphorylation of the AMP-activated protein kinase Snf1. Proteins were extracted from exponentially growing cells and were probed for Snf1, Snf1 phosphorylation (Snf1-P) and actin using AMPKAlpha, Phospho-AMPKAlpha (Thr172) and anti-actin antibodies.

To check whether Snf1 might act as a sensor of ATP decline under hypoxia in *C. albicans*, Snf1 phosphorylation was assessed in both normoxia and hypoxia with glucose and sucrose as carbon sources. Under normoxia, Snf1 was highly phosphorylated in WT cells that grew on sucrose as compared to cells utilizing glucose (**Figure 8B**). The same trend was observed for the *snf5* mutant. This suggests that sucrose is sensed as a non-optimal carbon sources regarding ATP generation as compared to glucose and that *snf5* is not compromised for the ability to sense the quality of the carbon sources. Under hypoxia, Snf1 phosphorylation remained unaffected in both WT and *snf5* cells for both sugars. This data indicate that Snf1 responds to carbon quality under normoxia while it is dispensable to sense the drop of ATP when cells experience hypoxia.

## Discussion

Inside the human host, fungal pathogens are exposed to a diverse spectrum of carbon sources that serve as a biosynthetic building blocks or are utilized to meet their energy demands. Although hypoxia is present in the human host at most foci of fungal infections, its contribution to fungal metabolism and fitness were underestimated so far. In this study, we have uncovered that Snf5, a subunit of SWI/SNF chromatin remodeling complex, is a major transcriptional regulator that links oxygen status to the metabolic capacity of *C. albicans*. Snf5 was required to activate genes of alternative carbon utilization and other carbohydrate related process specifically under hypoxia. Furthermore, *snf5* mutant exhibited an altered metabolome and lipidome reflecting that SWI/SNF chromatin remodelling activity plays an essential role in maintaining metabolic homeostasis and carbon flux in *C. albicans* under hypoxia. Snf5 was also required for both commensal growth in the gut and for systemic infection suggesting that the transcriptional control of metabolic flexibility under hypoxic environments is crucial for fungal fitness in the host (**Figure 9**).

**Figure 9.**
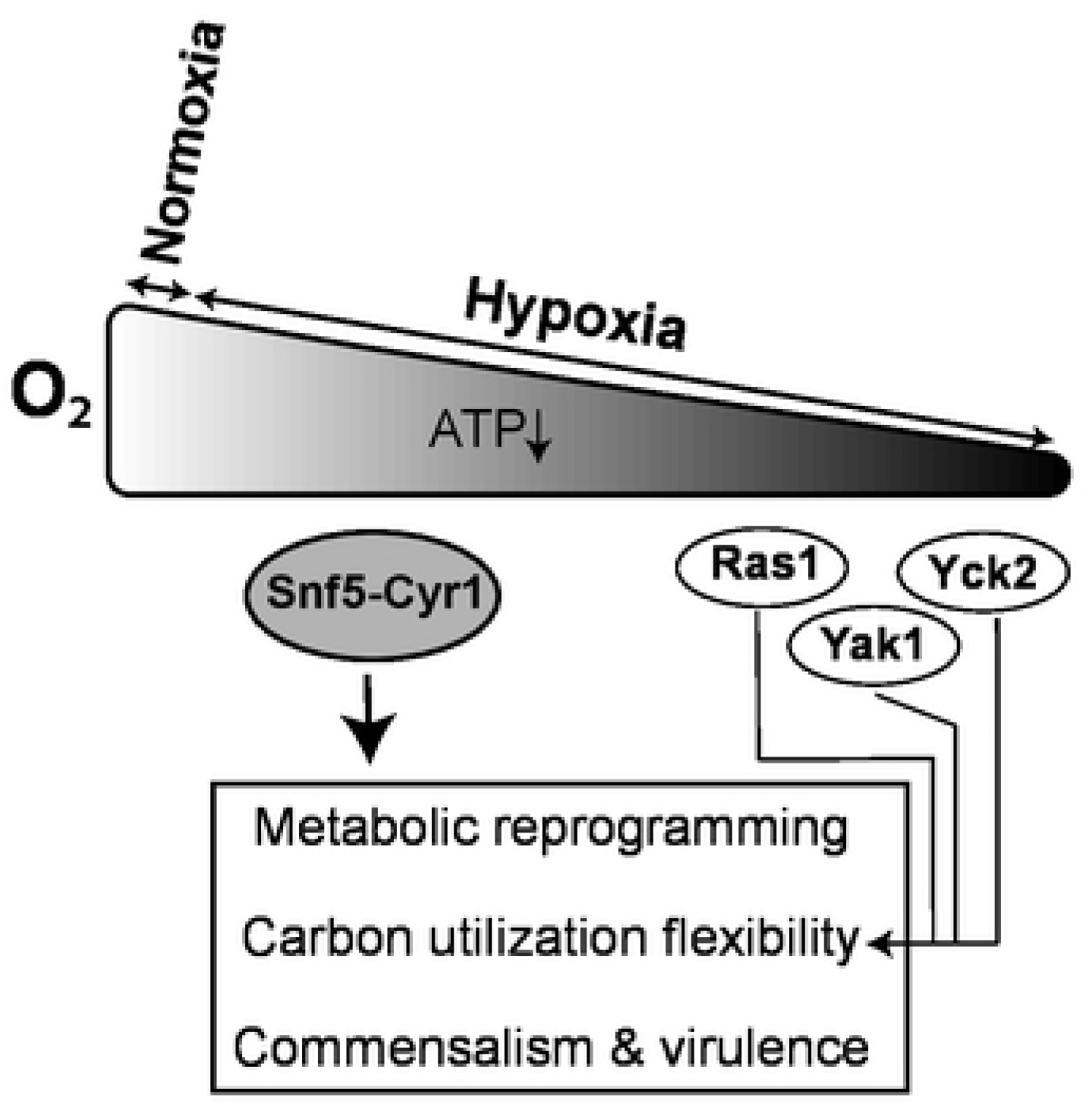
Summary of carbon metabolic flexibility control in *C. albicans* under hypoxia.

The ability of a microbial pathogen to thrive in human host is intimately linked to its ability to utilize a large spectrum of metabolites. In opposite to the saprophytic yeast *S. cerevisiae*, *C. albicans* is able to activate glycolytic, gluconeogenesis and glyoxylate cycles enzymes simultaneously to utilize both glucose and alternative carbon sources [13,55]. This evolutionary advantage might predispose *C. albicans* to proliferate in diverse niches with contrasting carbon sources inside the host. Intriguingly, genetic inactivation of Snf5 or other SWI/SNF subunits (Snf6, Snf2) had no impact on the fitness of *S. cerevisiae* in either normoxic or hypoxic environment while in the opportunistic yeast *C. glabrata,* deletion of Snf6 led to a similar growth defect as its homolog in *C. albicans*. As the role of SWI/SNF complex in metabolic adaptation seems to be specific to pathogenic yeasts, it is tempting to speculate that chromatin remodeling activity is an evolutionary driving force that might contribute to an enhanced fungal fitness [56]. Alternatively, the difference regarding the function of SWI/SNF in opportunistic *versus* saprophytic yeasts could be attributed to the fact that the function of a shared transcription factor, yet to be determined, that recruits SWI/SNF has rewired to accommodate host environments [57]. The specific role of SWI/SNF complex in controlling essential processes associated with fungal virulence makes it an interesting target for the development of antifungal therapeutics. While SWI/SNF complex is well conserved in eukaryotes including humans, some subunits such as Snf6, Taf14, Arp7 and Snf11 are unique to fungi and can be specifically targeted [41].

Our investigation uncovered that genetic inactivation of the different subunits of *C. albicans* SWI/SNF complex did not lead to the same phenotypic readouts. It is intriguing that the ATPase catalytic subunit Snf2 was dispensable for carbon metabolic flexibility under hypoxia while other subunits including Snf5, Snf6 and Swi1 were essential. A similar trend was also observed in *S. cerevisiae* where, for instance, mutants of only three SWI/SNF subunits (*snf2*, *swi3*, *taf14*) were required for growth in the presence of cell wall perturbing agents [58]. This differential requirement of SWI/SNF subunits corroborates the recent findings demonstrating a structural and functional modularity of the *S. cerevisiae* SWI/SNF complex [59,60]. In the budding yeast, SWI/SNF exhibited a modular architecture that reflect perfectly the contribution of each sub-module in transcriptional regulation of a specific subset of target genes [59]. In accordance with this, mutants of SWI/SNF subunits that exhibited metabolic flexibility defect in our study such as *snf6* and *snf5* or *swi1* and *arp9* belong to the same functional sub-complexes in the budding yeast. Alternatively, differential requirement of SWI/SNF subunit in *C. albicans* could be explained by the fact that deleting certain subunits might led to a non-functional aberrant complex that remains attached to its target promoters [58] and prevents redundant chromatin remodelling complex such as RSC to bind DNA and consequently compensates loss of SWI/SNF chromatin remodelling activity [49,50,51]. Accordingly, in *S. cerevisiae*, loss of Snf5 did not affect Snf2 occupancy to gene promoters and resulted in an aberrant SWI/SNF complex that functions as a dominant negative mutation by blocking recruitment of other redundant compensating chromatin remodelers [60].

In the budding yeast, recruitment of SWI/SNF by transcriptional factors such as Gal4, Hap4 and Gcn4 is important for transcriptional control at promoter regions [63,64]. To identify potential transcriptional regulators that require the chromatin remodelling activity of SWI/SNF to control carbon source utilization under hypoxia in *C. albicans*, we first interrogate genetic interactions between *SNF5* and *TYE7*, *MIG1*, *GAL4*, *ACE2*, *RTG1* and *RTG3* which are key transcription factors controlling glycolytic and other carbohydrate-related metabolic genes by dosage suppression (data not shown). Overexpressing of the candidate regulators in *snf5* did not result in any phenotypic enhancement or genetic epistasis suggesting that Snf5 control metabolic flexibility independently of those characterized transcriptional regulators. A similar finding was obtained when increasing the dosage of *SNF5* in mutants of the aforementioned transcription factors or when using CHI. This suggests that our work has uncovered a unique regulatory circuit that controls carbon metabolism in response to oxygen levels. Among candidates of signaling pathways in which Snf5 might operates, phenotypic analysis revealed that mutants of Ras1-cAMP-PKA pathway, as well as mutant of Yak1 and Yck2 kinases exhibited a similar carbon flexibility phenotype as did *snf5* under hypoxia. Our CHI analysis indicates a functional link between Snf5 and the adenylyl cyclase Cyr1 and we also uncovered novel independent role of Ras1, Yck2 and Yak1 in metabolic adaptation in response to hypoxia. Future efforts will lend further insights into mechanisms by which these regulators transmit oxygen status to the fungal metabolic machinery.

A critical but still poorly understood aspect of eukaryotic biology is how cells communicate the status of oxygen to the metabolic machinery. As oxygen becomes limiting for oxidative phosphorylation, a direct consequence of hypoxia is the shift from respiration to fermentation mode to generate ATP. We found that ATP levels decreased significantly when *C. albicans* experienced hypoxia which might prime metabolic adaptation to compensate such energy failure. In metazoan, hypoxia led to a reduction of ATP levels and the activation of the Snf1/AMPK protein kinase to maintain energy homeostasis [1]. Our data showed that exposure of *C. albicans* cells to hypoxia did not led to Snf1 activation suggesting that Snf1 is dispensable to maintain ATP homeostasis in oxygen-limiting environments. This begs the question of how *C. albicans* senses and signals ATP depletion under hypoxia. Snf2 is the catalytic subunit of SWI/SNF complex that couples ATP hydrolysis through its ATPase activity to nucleosome repositioning and histone eviction [65]. Consequently, SWI/SNF activity should be sensitive to cellular fluctuation of ATP levels and suggests that this chromatin remodelling complex might directly link energy status in the cell to gene expression regulation. While the role of ATP sensing was not attributed to SWI/SNF complex in either microbial eukaryotes or metazoans so far, several observations support this hypothesis. Our data indicated that *C. albicans* SWI/SNF govern the transcriptional control of the same category of genes that are modulated to maintain ATP homeostasis in eukaryotic cells [53,66,67]. In *S. cerevisiae* and mammalian cells, the low-energy checkpoint Snf1/AMPK induces energy generating (beta-oxidation, carbon utilization, carnitine metabolism) and represses energy consuming reactions (fatty acid, lipid and protein biosynthesis). Our transcriptional analysis in *C. albicans* uncovered that Snf5 was required for the activation of replenishing ATP pathways including alterative carbon metabolism and fatty acid beta-oxidation, and the down regulation of ATP consuming processes of macromolecule biosynthesis including proteins and lipids (**Table 1 and S4**). This was also recapitulated in *snf5* metabolome with a significant increase of different classes of lipids and a decrease of CoA and Acetyl-CoA. This indicates that *C. albicans* SWI/SNF is mimicking the role of Snf1/AMPK as energy gauge under hypoxia. Alternatively, Cyr1 might also act as an ATP sensor to reprogram cellular metabolism in response to hypoxia as this protein has an ATP binding domain. A similar role was proposed to this adenylate cyclase in the regulation of Ras1-modulated virulence pathways as a consequence of ATP depletion [68].

In conclusion, this study yielded unprecedented insight into the regulatory circuit that control carbon metabolism in response to oxygen levels. Snf5 subunit of the SWI/SNF chromatin remodelling complex provides a nexus for integrating oxygen status to the carbon metabolic machinery and fungal fitness. As hypoxia is the predominant condition inside the host and since *C. albicans* utilizes different alternative carbon sources when persisting as a commensal [69] or infecting its host [70,71], SWI/SNF represents thus an attractive target for antifungal therapy.

## Material and methods

### Ethics statement

All procedures used to study intestinal colonization in mice were approved by the Tufts University Institutional Animal Care and Use Committee (approved protocol B2015-147). Mice were sacrificed by CO_2_ inhalation following a method of euthanasia that is conform to the recommendations from the panel on Euthanasia of the American Veterinary Medical Association.

### Strains, mutant collections and growth conditions

*C. albicans* strains were cultured at 30°C in yeast-peptone-dextrose (YPD) medium supplemented with uridine (2 % Bacto peptone, 1 % yeast extract, 2 % w/v dextrose, and 50 mg/l uridine). Alternative and non-fermentable carbon sources (fructose, sucrose, galactose, mannose, maltose, oleic acid, glycerol, acetate, lactate, ethanol, mannitol and sorbitol) were used at 2 % w/v. WT and mutant strains used in this study together with diagnostic PCR primers are listed in **Table S7.** The transcriptional factor [33] mutant collection used for metabolic flexibility screens were acquired from the genetic stock center (http://www.fgsc.net). The transcriptional regulator [32] mutant collection was kindly provided by Dr. Dominique Sanglard (University of Lausanne).

For filamentation assays, cells were grown at 37°C in YPD supplemented with either 10% fetal bovine serum (FBS; Invitrogen) or 2.5 mM N-acetyl-D-glucosamine (GlcNAc; Sigma) or in Spider medium [72]. For growth under hypoxic conditions, cells were spotted on YPS (2% Bacto peptone, 1% yeast extract, 2% sucrose, 2% agar) plates and incubated in an anaerobic chamber (Oxoid; HP0011A) at 37°C. The chamber was flushed daily with nitrogen to remove oxygen and any by-products. For agar invasion assay, 4 µl of OD_600_ of 2 of exponentially *C. albicans* cells was seeded on YPD agar plate and incubated for 3 days at 30 °C. The plates were photographed before and after being washed with phosphate-buffered saline (PBS). Furthermore, and for each *C. albicans* strain, agar sections were performed and photographed.

### Genetic screen and growth assays

For each mutant collection, strains were arrayed using a sterilized 96-well pin tool on Nunc Omni Trays containing either YPD-agar, YPS-agar or YPG-agar and colonies were grown for four days at 30°C under normoxic (21% O_2_) or hypoxic conditions (5% O_2_). Plate were then imaged using the SP-imager system. A growth score was given for each mutant (**Table S1**). Each mutant hit was confirmed individually using dilution spot assay. *snf5*/*SNF5* mutant was subjected to epistatic analysis using the concept of complex haploinsufficiency (CHI) [48,73] with deletions of one allele of *RAS1*, *YCK2*, *YAK1* and *CYR1*. Gene deletion was performed as previously described [74]. The complete set of primers used to generate deletion cassettes and to confirm gene deletions are listed in **Table S7**.

### Expression analysis by Microarrays and qPCR

Overnight cultures of *snf5* mutant and WT strains were diluted to an OD_600_ of 0.1 in 100 ml of fresh YPS-uridine medium, grown at 30°C to an OD_600_ of 0.4. Cells were harvested by centrifugation at 3,000 x g for 5 min, and the pellet was washed with PBS and resuspended in 400 µl of YPS medium. Half of the *C. albicans* cell suspension was used to inoculate aerated flasks containing fresh YPS medium (normoxia), and the second half was added to bottles containing fresh YPS flushed with nitrogen to remove oxygen (hypoxia). Candida cells were then incubated at 30°C under agitation at 220 rpm for 1 hour. Cells were then centrifuged for 2 min at 3,500 rpm, the supernatants were removed, and the samples were quick-frozen and stored at −80 °C. Total RNA was extracted using an RNAeasy purification kit (Qiagen) and glass bead lysis in a Biospec Mini 24 bead-beater. Total RNA was eluted, assessed for integrity on an Agilent 2100 Bioanalyzer prior to cDNA labeling, microarray hybridization and analysis [75]. The GSEA Pre-Ranked tool (http://www.broadinstitute.org/gsea/) was used to determine statistical significance of correlations between the transcriptome of the *snf5* mutant with a ranked gene list or GO biological process terms as described by Sellam *et al*. [24]. Differentially expressed transcripts in **Table S1** were identified using Welch’s t-test with a false-discovery rate (FDR) of 5% and 1.5-fold enrichment cut-off.

For qPCR experiments, cell cultures and RNA extractions were performed as described for the microarray experiment. cDNA was synthesized from 1µg of total RNA using High-Capacity cDNA Reverse Transcription kit (Applied Biosystems). The mixture was incubated at 25°C for 10 min, 37°C for 120 min and 85°C for 5 min. 2U/µl of RNAse H (NEB) was added to remove RNA and samples were incubated at 37°C for 20 min. qPCR was performed using a LightCycler 480 Instrument (Roche Life Science) for 40 amplification cycles with the PowerUp™ SYBR® Green master mix (Applied Biosystems). The reactions were incubated at 50°C for 2 min, 95°C for 2min and cycled for 40 times at 95°C, 15 s; 54°C, 30 s; 72°C, 1 min. Fold-enrichment of each tested transcripts was estimated using the comparative ΔΔCt method. To evaluate the gene expression level, the results were normalized using Ct values obtained from Actin (*ACT1*, C1_13700W_A). Primer sequences used for this analysis are summarized in Supplemental **Table S7.**

### Galleria virulence assay

For *Galleria mellonella* studies, overnight cultures were washed twice and resuspended in PBS then cell number was determined by Coulter counter. *G. mellonella* larvae weighing 180 ± 10 mg were injected between the third pair prothoracic legs with 5 µl of suspension (10^6^ cells). Infected larvae were incubated at 37°C. Four replicates, each consisting of 40 larvae, were carried out with survival rates measured daily for a period of 6 days. Death was determined based on the lack of response to touch and the inability to right themselves. Kaplan-Meier survival curves were created and compared with the log-rank test (GraphPad Prism 5).

### Intestinal colonization assay

Female Swiss Webster mice (18–20 g) were treated with streptomycin (2 g/l), gentamycin (0.1 g/l), and tetracycline (2 g/l) in their drinking water throughout the experiment beginning 4 days prior to inoculation. Mice were inoculated with *C. albicans* by oral gavage (5×10^7^ *C. albicans* cells in 0.1 ml), as described previously [76]. Colonization was monitored by collecting fecal pellets (produced within 10 minutes prior to collection) at various days post-inoculation, homogenizing in PBS, plating homogenates on YPD agar medium supplemented with 50 µg/ml ampicillin and 100 µg/mL streptomycin. Mice were sacrificed by CO_2_ inhalation with a regulated flow of CO_2_ on day 10 post-inoculation. *C. albicans* concentrations in stomach, cecum and colon were measured by plating as described above. As the *snf5* strain clumps, qPCR was also used to quantify *C. albicans* concentration in cecum and compared to plating. We found a good correlation between these two methods for quantification (*R*² = 0,9122). Composite results of plating from two experiments were shown.

### Adhesion assay

The ability of *C. albicans* strains to adhere to host cells was assessed on the human colon epithelial cell line HT-29 (ATCC; HTB-38). HT-29 cells were plated in 12-well plate to obtain 100% confluent cells. Overnight cultures of *C. albicans* strains were wash twice with PBS and cellular concentration was adjusted to 12×10^6^ cells/ml in McCoy’s medium. HT-29 cells medium was removed and 6×10^6^ cells of yeast strain was added per well. A co-incubation was performed at 37°C and 5% CO_2_ for 1 hour. Non-adherent cells were removed by rinsing five times with 1 ml PBS and cells were then fixed with 4% paraformaldehyde. HT-29 cells were permeabilized with 0.5% Triton X-100. Next adherent fungal cells were stained with 2 µM calcofluor white during 30 min in the dark at room temperature. Adherent cells were visualised using Cytation 5 high-content microscope with 20x magnification and DAPI filter. For each well, 20 fields were at least photographed.

### HT29 and J774A.1 damage assay

Damage to the human colon epithelial cell line HT-29 and macrophages J774A.1 (ATCC TIB-67) were assessed using a lactate dehydrogenase (LDH) cytotoxicity detection kit^PLUS^ (Roche), which measures the release of LDH in the growth medium. The manufacturer’s protocol was followed. HT-29 cells were grown in 96-well plate as monolayers in McCoy’s medium supplemented with 10% FBS at 1×10^4^ cells per well, and J774A.1 were grown in DMEM medium supplemented with 10% FBS at 1.5×10^4^ cells per well in a 96-well plate and incubated at 37°C with 5% CO_2_ overnight. HT-29 and J774A.1 cells were then infected with *C. albicans* cells at MOI cell:yeast of 1:2 for 24 h at 37°C with 5% CO_2_. Following incubation, 100 µl of supernatant was removed from each experimental well and LDH activity in this supernatant was determined by measuring the absorbance at 490 nm (OD_490_).

### ATP quantification

*C. albicans* cultures were grown overnight in YPD medium and were washed twice with PBS. Cells were then resuspended in either YPD, YPG or YPS to an OD_600_ of 0.5. A total of 90 µl of these cell suspensions were transferred to a white-opaque 96-well plate. Cells were then incubated in either hypoxia (5% oxygen) or normoxia (21% oxygen) for 3 hours. An equal amount of BacTiter-Glo reagent was added to each well as described by the manufacturer (BacTiter-Glo Microbial Cell Viability, Promega). For ATP quantification under normoxia, the plate was mixed for 5 min on an orbital shaker and incubated 15 minutes at room temperature. For hypoxic conditions, plate with yeast suspension was incubated 15 minutes at 30°C with 5% oxygen.

### Metabolomic analysis

Metabolomics analysis was carried out in collaboration with Metabolon (Durham, NC, USA). Cell cultures were performed as described for the microarray experiment. A total of five biological replicates were submitted to Metabolon for metabolite profiling. To remove protein, dissociate small molecules bound to protein, and to recover chemically diverse metabolites, proteins were precipitated with methanol under vigorous shaking for 2 min followed by centrifugation. The resulting extracts were analysed by GC/MS and LC/MS. All metabolites with mean values that had significant differences (p<0.05) between treated and untreated samples were considered as enriched (>1.5-fold) or depleted (<1.5-fold).

### Western blot analysis

A total of 5 ml of YPD overnight cultures of WT and *snf5* mutant strains were washed twice with sterile water and then diluted to an OD_600_ of 2 in 20 ml of YPD or YPS under normoxic and hypoxic conditions and incubated for 3 hours at 30°C. Cells were collected by centrifugation and frozen in liquid nitrogen. Cell pellets were thawed on ice, washed with sterile water, and 2 ml of YeastBuster™ (Millipore Sigma), supplemented with cOmplete™ EDTA-free Protease Inhibitor Cocktail (sigma) and PhosStop Phosphatase Inhibitor Cocktail (Roche), was added. The lysis was performed at room temperature and the insoluble cell debris were removed by centrifugation at 16,000 × g for 20 min. Protein concentration was quantified with RC-DC protein assay II (Biorad). An equal amount of 40 µg of proteins was separated on an 8% SDS polyacrylamide gel and transferred onto nitrocellulose membranes. The membrane was blocked using the StartingBlock™ Blocking Buffer (Thermofisher) during 15 min at room temperature. To detect Snf1 phosphorylation the Phospho-AMPKα (Thr172) Antibody (Cell Signaling #2531) was diluted to 1:1000 in TBST 5% BSA and incubated overnight at 4°C. Membranes were washed four times in TBST and then incubated with anti-rabbit HRP antibody diluted to 1:5000 for one hour at room temperature. Membranes were washed four times with TBST and 2 times with TBS and signals were detected with Pierce ECL Western Blotting Substrate (Thermo Scientific). For Snf1 and actin detections, AMPKα (Cell Signaling #2532) and actin (Genscript, A00730-200) antibodies were diluted to 1:1000 in TBST 5% BSA and incubated overnight at 4°C following by the same procedure as describe previously.

## Acknowledgments

We are grateful to Aaron Mitchell (Carnegie Mellon University), Dominique Sanglard (Le Centre Hospitalier Universitaire Vaudois (CHUV)-Université Lausanne), Doreen Harcus (National Research Council Canada), Karl Kuchler (Medical University of Vienna) and Guilhem Janbon (Institut Pasteur-Paris) for providing strains.

## Supporting information

**Figure S1.** (**A**) Growth defect of the SWI/SNF subunit mutants *swi1*, *snf6* and *arp9* in alternative carbon sources under hypoxia. Mutants were form the GRACE collection and were grown under repressing conditions (100 µg/ml tetracycline) for 4 days at 30°C. (**B-C**) Metabolic flexibility phenotype of different SWI/SNF subunit mutants of *S. cerevisiae* (**B**) and the opportunistic yeast *C. glabrata* (**C**). Growth of the WT strain of *S. cerevisiae* (BY4741) and *C. glabrata* (HTL) and the SWI/SNF mutants in media with the indicated carbon sources under both normoxic (21% O_2_) and hypoxic (5% O_2_) are shown.

**Figure S2.** (**A**) qPCR validation of altered expression levels of *GLK1*, *MAL32*, *MLS1*, *PFK1*, *ALD6*, *MDH1* and *FBP1* in both WT and *snf5* mutant strains under hypoxia. Relative expression levels of the seven transcripts were assessed by real-time qPCR and normalized to *ACT1* relative to normoxic conditions. Values are the mean from at least two independent experiments. (**B**) Venn diagram showing overlaps between genes differentially regulated in *snf5* mutant and promoters bound by Snf6 as shown by Tebbji *et al.* [77]. Relevant functional categories are shown. **(C)** Transcript level of the transcription factor *TYE7* in both WT and *snf5* mutant strains under hypoxia relative to normoxic conditions.

**Table S1.** Raw data of the genetic survey for transcriptional regulators required for metabolic adaptation in different carbon sources under low oxygen concentration

**Table S2.** Transcripts differentially expressed in *snf5* mutant using a 1.5-fold change cut-off and a 5% false discovery rate.

**Table S3.** Raw data of the WT and *snf5* mutant strains.

**Table S4.** Gene Set Enrichment Analysis (GSEA) of *snf5* mutant transcriptome under hypoxia.

**Table S5.** Lists of statistically enriched or depleted metabolites in *snf5* mutant under both normoxia and hypoxia as compared to the WT strain as presented in Venn diagrams of **Figure 4B**.

**Table S6.** Full metabolomic data of *snf5* mutant under both normoxia and hypoxia as compared to the WT strain.

**Table S7.** List of *C. albicans* strains and primers used in this study.

